# Patient-derived *phosphoribosyl pyrophosphate synthetase* mutations in *Drosophila* result in autophagy and lysosome dysfunction

**DOI:** 10.1101/438747

**Authors:** Keemo Delos Santos, Christine Yergeau, Nam-Sung Moon

## Abstract

Phosphoribosyl pyrophosphate synthetase (PRPS) is a rate-limiting enzyme in nucleotide metabolism. While missense mutations of *PRPS1* have been identified in neurological disorders such as Arts syndrome, little is known on how they contribute to pathogenesis. We engineered *Drosophila PRPS* (*dPRPS*) alleles that carry patient-derived PRPS missense mutations. Although *dPRPS* mutant flies develop normally, they have profound defects in autophagy induction and lysosome function. Consequently, *dPRPS* flies are sensitive to nutrient deprivation as they are unable to break down lipid storage by macroautophagy. In addition, we provide evidence showing that *dRPPS* is required for proper cellular response to oxidative stress, providing a possible mechanism by which PRPS1 dysfunction contributes to neurological disorders.

## Introduction

Phosphoribosyl pyrophosphate synthetase (PRPS) is a rate-limiting enzyme in the biosynthesis of purine, pyrimidine and pyridine nucleotides (Fig. 1A). Purine and pyrimidine nucleotides are the building blocks of RNA and DNA while pyridine nucleotides, such as NAD and NADP, are important co-factors in many enzymatic reactions. PRPS produces phosphoribosyl pyrophosphate (PRPP), a common precursor of the five-carbon sugar subunit of nucleotides (Hove-Jensen et al. 2017). The essential role of PRPS in nucleotide metabolism is illustrated through the conservation of this enzyme among all free-living organisms ranging from *E.coli* to humans.

**Figure 1.**
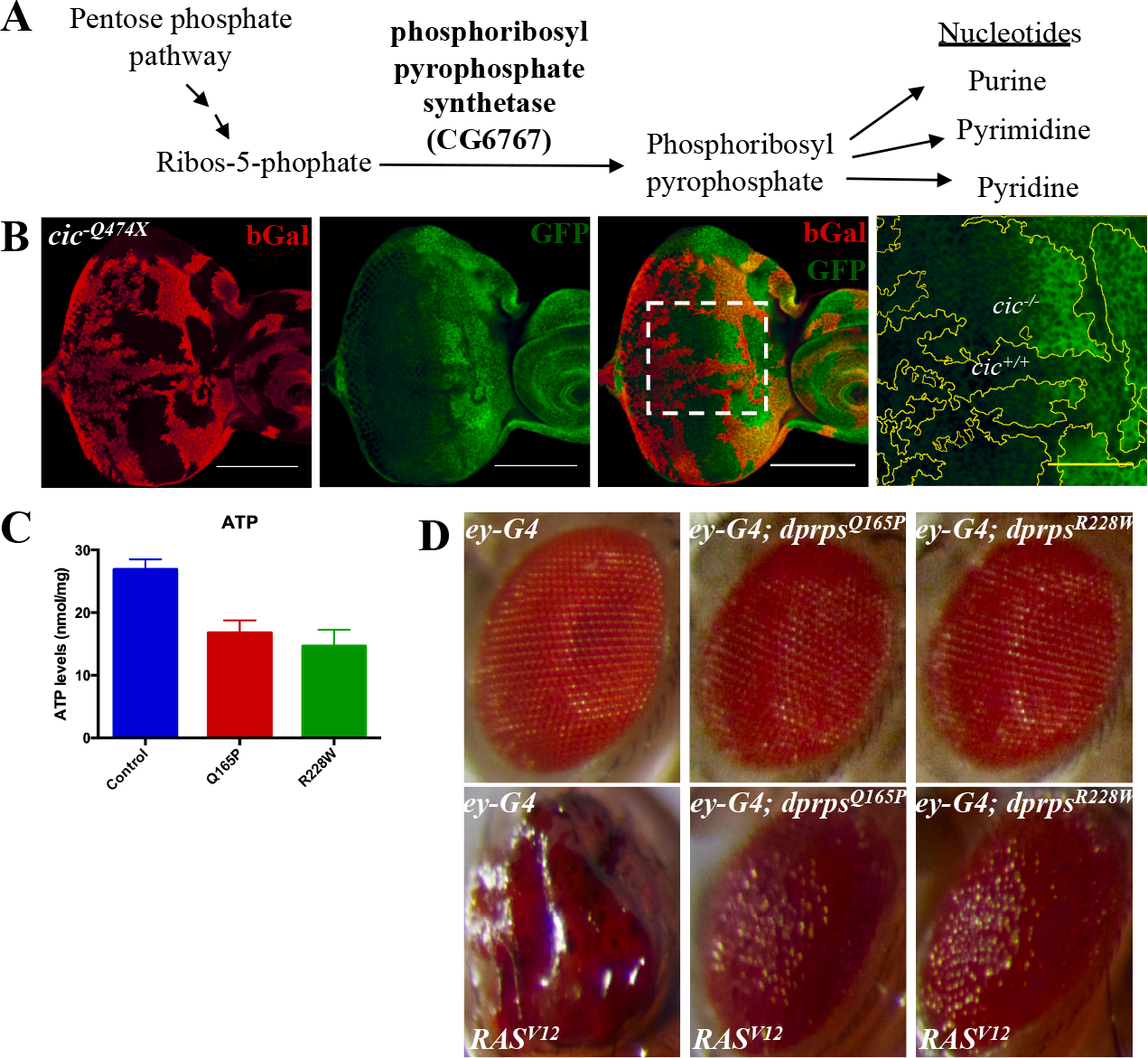
*phosphoribosyl pyrophosphate synthetase* (*PRPS*) is a Capicua-regulated gene that is required for the Ras-induced tumor phenotype. (A) Schematic of the role of PRPS in nucleotide biosynthesis is shown. PRPS converts ribose-5-phosphate from the pentose phosphate pathway into phosphoribosyl pyrophosphate (PRPP), which is required for synthesis of various nucleotides. (B) Somatic clones of *capicua* (*cic*) mutant cells are generated in third instar larval eye imaginal discs. *cic* mutant clones are marked by the lack of mGal signal (red). A GFP protein trap allele of *dPRPS* (green) was used to monitor expression of PRPS in control and *cic* mutant clones. A magnified view of the indicated area (white box) is also presented. Outlined in yellow denote the border between wild type and *cic* mutant clones. Scale bars: 100 μm for white and 30 μm for yellow. (C) PRPS mutant alleles carrying patient-derived missense mutations were generated using the *Drosophila* CRISPR/Cas9 system (*dPRPS*^*Q165P*^ *and dPRPS*^*R228W*^; see materials and methods). The graph shows the relative ATP levels in control and *dPRPS*^*Q165P*^ and *dPRPS*^*R228W*^ adult females. Error bars indicate standard deviation from triplicated experiments (D) The effect of overexpressing an activated form of Ras (*Ras*^*V12*^) in control (left panel), *dPRPS*^*Q165P*^ (middle panel) and *dPRPS*^*R228W*^ (right panel) eyes are shown.

In humans, three *PRPS* orthologs exists, *PRPS1* to *3*. Notably, *PRPS1* mutations were identified in a number of neurological disorders: Arts syndrome, Charcot-Marie-Tooth disease, and nonsyndromic sensorineural deafness (de Brouwer et al. 2007; Al-Maawali et al. 2015). All *PRPS1* mutations identified from patients are missense mutations affecting enzymatic activity to varying degrees. The fact that no nonsense mutations were identified suggests that *PRPS1* is essential for embryonic development and patients with missense mutations retain a certain level of PRPS1 function. Thus far, PRPS orthologs have been knocked out in a number of animal models. *PRPS1* knockout mice were generated as a part of a high-throughput screening of mouse genes important for skeletal phenotypes (Brommage et al. 2014). Not surprisingly, *PRPS1* was identified as an X-linked gene that is required for animal viability. In contrast, *PRPS2* knockout mice are viable and fertile with no discernable developmental defects, suggesting that other mouse *PRPS* orthologs compensate for the loss of *PRPS2* (Cunningham et al. 2014). Interestingly, while *PRPS2* is non-essential for development, *PRPS2* knockout mice are resistant to *Eμ–Myc*-driven cancer development, suggesting that PRPS2 is specifically required for tumorigenesis (Cunningham et al. 2014). A zebrafish model of PRPS deficiency has also been recently generated (Pei et al. 2016). The *prps1a* and *prps1b* double mutant animals fail to properly develop but show some phenotypic similarity to human PRPS1-associated diseases. Importantly, only the null alleles of *PRPS* were generated in both mouse and zebrafish models and the biological consequence of patient-derived *PRPS1* mutations have not been directly tested.

*Drosophila* has only one PRPS ortholog (dPRPS) coded by an uncharacterized gene *CG6767*. dPRPS has 89% protein sequence identity with the mammalian PRPS1 (Supplemental Fig. S1A). To investigate the *in vivo* function of dPRPS, we generated *dPRPS* alleles that carry mutations identified from Arts syndrome, *dPRPS*^*Q165P*^ and *dPRPS*^*R228W*^ (Supplemental Fig. S1A and S1D, and (de Brouwer et al. 2010; Al-Maawali et al. 2015). Interestingly, while *dPRPS*^*Q165P*^ and *dPRPS*^*R228W*^ flies are viable and fertile, they are highly sensitive to nutrient withdrawal. We discovered that their susceptibility to starvation is, at least in part, caused by their failure to mobilize their lipid reserves due to profound defects in autophagy and lysosome function. Further analysis also revealed that dPRPS plays a critical function during cellular response to oxidative stress, providing a possible explanation by which PRPS1 dysfunction promotes neurological disorders.

## Results

### Patient-derived *PRPS* mutations suppress a Ras-induced hyperplastic phenotype

Capicua (Cic) is a transcriptional repressor downstream of the EGFR/Ras pathway that regulate various cellular processes (Simón-Carrasco et al. 2018). We and others have previously identified Cic as an important determinant of cellular proliferation (Tseng et al. 2007; Krivy et al. 2012). A recent study using DNA adenine methyltransferase identification (DamID) technique identified *CG6767* (dPRPS) as one of the Cic targets (Supplemental Fig. S1A and Jin et al. 2015). To confirm this finding, we tested if *CG6767* expression is regulated by Cic using a protein trap allele, *GFP-trap* (Supplemental Fig. S1B). Comparing the GFP expression levels in eye imaginal discs containing *cic* mutant clones (*cic*^*Q474X*^) confirmed that dPRPS is normally repressed by Cic (Fig. 1B). Because the publicly available *dPRPS* alleles are early larval lethal, it was difficult to use them to investigate *in vivo* function of dPRPS (Supplemental Fig. S1C). Given the high sequence similarity and that patient-derived *PRPS1* missense mutations likely represent hypomorphic mutations, we engineered *dPRPS* alleles that carry the mutations identified from Arts syndrome via CRISPR/Cas9, *dPRPS*^*Q165P*^ and *dPRPS*^*R228W*^ (Supplemental Fig. S1A and S1D). Interestingly, *dPRPS*^*Q165P*^ and *dPRPS*^*R228W*^ flies are viable and fertile with no discernable developmental defects, suggesting that they retain a sufficient amount of PRPS function for animal development. To confirm that dPRPS function is indeed compromised in *dPRPS*^*Q165P*^ and *dPRPS*^*R228W*^ flies, two tests were performed. First, we determined the relative ATP levels in control and *dPRPS* mutant flies. The ATP levels were significantly reduced in *dPRPS* mutant flies, suggesting that the enzymatic activity is impaired (Fig. 1C). Second, we tested if *dPRPS*^*Q165P*^ and *dPRPS*^*R228W*^ mutations can suppress the Ras-induced tumor phenotype. Since Cic is a crucial factor regulating proliferation downstream of the EGFR/Ras pathway, we reasoned that Ras may require dPRPS to promote hyperproliferation. Overexpressing an activated form of Ras (Ras^V12^) in the *Drosophila* eye produces hyperplastic overgrowth (Karim and Rubin 1998). Strikingly, the Ras^V12^-induced phenotype is strongly suppressed in the *dPRPS*^*Q165P*^ or *dPRPS*^*R228W*^ mutant background, indicating that dPRPS is required for Ras-dependent hyperplastic overgrowth (Fig. 1D). Taken together, our results suggest that *dPRPS*^*Q165P*^ and *dPRPS*^*R228W*^ are hypomorphic alleles of *dPRPS* that maintain a sufficient amount of PRPS function to support animal development.

### *dPRPS* mutants have lipid mobilization defects

The *Drosophila* fat body functions as an energy reserve in the form of lipid droplets and plays a role similar to the mammalian liver and adipose tissue (Zhang and Xi 2015). Interestingly, publicly available expression data from large-scale microarrays and RNA-sequencing analyses indicate that *dPRPS* is highly expressed in the *Drosophila* fat body (Supplemental Fig. S1E). This was confirmed by the *dPRPS GFP-trap* allele that showed strong expression (Supplemental Fig. S1F) and led us to hypothesize that dPRPS plays an important function in the fat body. Because animal survival during starvation is tightly linked to fat usage (Zhang and Xi 2015), we examined the sensitivity of *PRPS^Q165P^* and *dPRPS*^*R228W*^ flies to starvation. Indeed, upon complete nutrition withdrawal, *dPRPS*^*Q165P*^ and *dPRPS*^*R228W*^ flies die about two days faster than age-matched control flies (Supplemental Fig. S2). We next visualized the lipid droplets in the fat body during the progression of starvation (Fig. 2A). While the size of lipid droplets decreases in control flies, it remains relatively unchanged in *dPRPS*^*Q165P*^ and *dPRPS*^*R228W*^ flies during starvation (Fig. 2A and 2B). Additionally, quantification of triglyceride (TAG) levels, the primary lipid form in which fat is stored, revealed a similar trend wherein TAG levels are significantly reduced in the control flies during starvation, but are relatively unchanged in the *dPRPS* mutants (Fig. 2C). This observation suggests that the increased sensitivity to starvation in *dPRPS*^*Q165P*^ and *dPRPS*^*R228W*^ flies may be caused by an inability to use their lipid reserves. To test this hypothesis, we overexpressed *Lipase 4* (*Lip4*), the primary lipase used in starvation-mediated lipolysis, in control and *dPRPS* fat bodies (Vihervaara and Puig 2008). *Lip4* overexpression had little to no effect on the survival of control flies during starvation (Fig. 2D). However, *Lip4* overexpression in *dPRPS*^*Q165P*^ and *dPRPS*^*R228W*^ flies significantly improved their survival, suggesting that defects in lipid mobilization contribute to their hypersensitivity to nutrient withdrawal. Notably, accumulation of lipid droplets in glia has been identified as one of the common features in *Drosophila* models of neurodegeneration (Liu et al. 2015). Given that human *PRPS1* is mutated in a number of neurological disorders, we examined if a similar phenotype can be observed in *dPRPS*^*Q165P*^ and *dPRPS*^*R228W*^ flies. Indeed, we observed lipid droplet accumulation in *dPRPS*^*Q165P*^ and *dPRPS*^*R228W*^ pupal eyes but not in control (Fig. 2E). This indicates that the lipid mobilization defect is not limited to the fat body during nutrient deprivation and may be a common consequence of dPRPS dysfunction.

**Figure 2.**
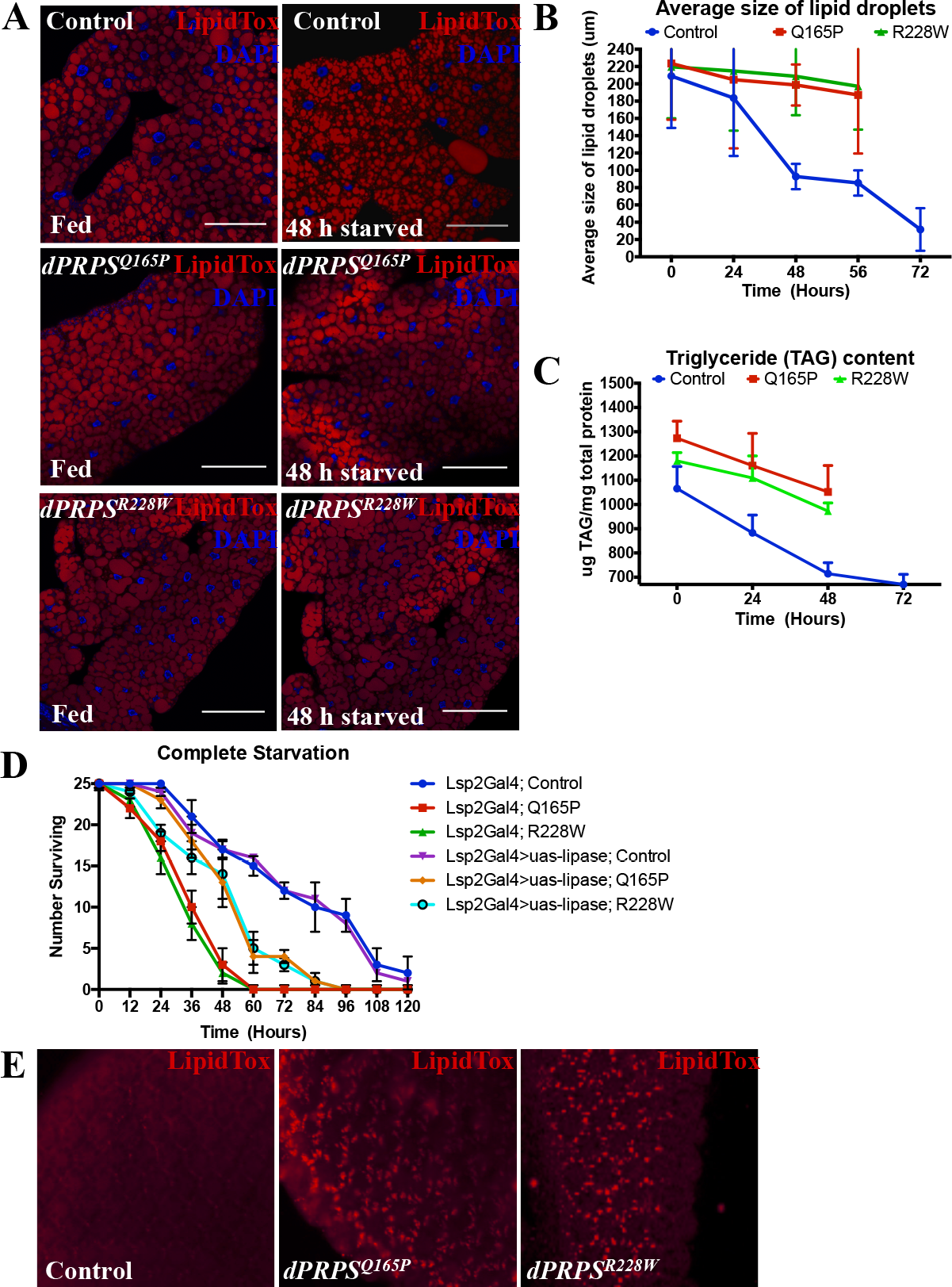
*dPRPS* mutants have lipid mobilization defects. (A) Adult fat bodies from control and *dPRPS* mutant flies under fed (left panel) and starved (right panel) conditions are stained with 4′,6-diamidino-2-phenylindole (DAPI, blue) and LipidTOX (red) to visualize nuclei and lipid droplets, respectively. Scale bar: 100μm. (B) A graph showing the size of lipid droplets in control, *dPRPS*^*Q165P*^ and *dPRPS*^*R228W*^ flies at indicated time points during starvation is presented. (C) A graph showing normalized triacylglycerol (TAG) levels in control and *dPRPS* mutant adult flies during starvation is presented. (D) The effect of expressing a lipase under the control of a fat body-specific driver, *lsp2-Gal4*, is shown. Error bars indicate standard deviation from triplicated experiments. (E) Pupal eye discs at 42 hours after pupa formation from control and *dPRPS* mutant flies stained with LipidTOX are shown.

### dPRPS is required for starvation-induced autophagy

During starvation, autophagy is one of the primary mechanisms to mobilize lipids (Scott et al. 2004). Intracellular lipids are sequestered in double membrane vesicles called autophagosomes and delivered to the lysosome for their eventual degradation into fatty acids for energy production or macromolecular synthesis (Lum et al. 2005; Mizushima 2007). To assess whether autophagy is deregulated in *dPRPS* mutants, we expressed GFP-Atg8, a commonly used molecular marker of autophagy, in the fat body (Nagy et al. 2015). Indeed, control fat bodies show a basal level of autophagy under fed conditions, which increases during starvation (Fig. 3A upper panel and 3C). Strikingly, *dPRPS* mutant flies have an almost undetectable level of autophagy under both fed and starved conditions (Fig. 3A lower panel and 3C). In addition, GFP-tagged Lamp1, a lysosomal protein, was used to monitor the expansion of the lysosome compartment during autophagy (Nagy et al. 2015). Similar to what was observed with GFP-Atg8, *dPRPS* mutant flies have almost an undetectable level of GFP-Lamp1 under fed and starved conditions. (Fig. 3B and 3D). To functionally confirm the autophagy defect revealed by the molecular markers, GFP-Atg8 and GFP-Lamp1, we examined the effect of depleting core autophagy proteins *atg8* and *atg16* in *dPRPS* fat bodies (Mizushima et al. 2011; Nagy et al. 2015). *atg8* or *atg16* depletion should have a minimal effect in *dPRPS* mutant flies if autophagy is already compromised as shown in Figure 3C and 3D. Indeed, while *atg8* or *atg16* knockdown decreased the survival rate of wild-type flies upon starvation, it had no significant effect on the survival of *dPRPS* mutant flies (Fig. 3E and 3F). Overall, these data suggest that dPRPS dysfunction caused by patient-derived mutations affects autophagy induction and may explain why *dPRPS* mutants are unable to breakdown lipid droplets during starvation.

**Figure 3.**
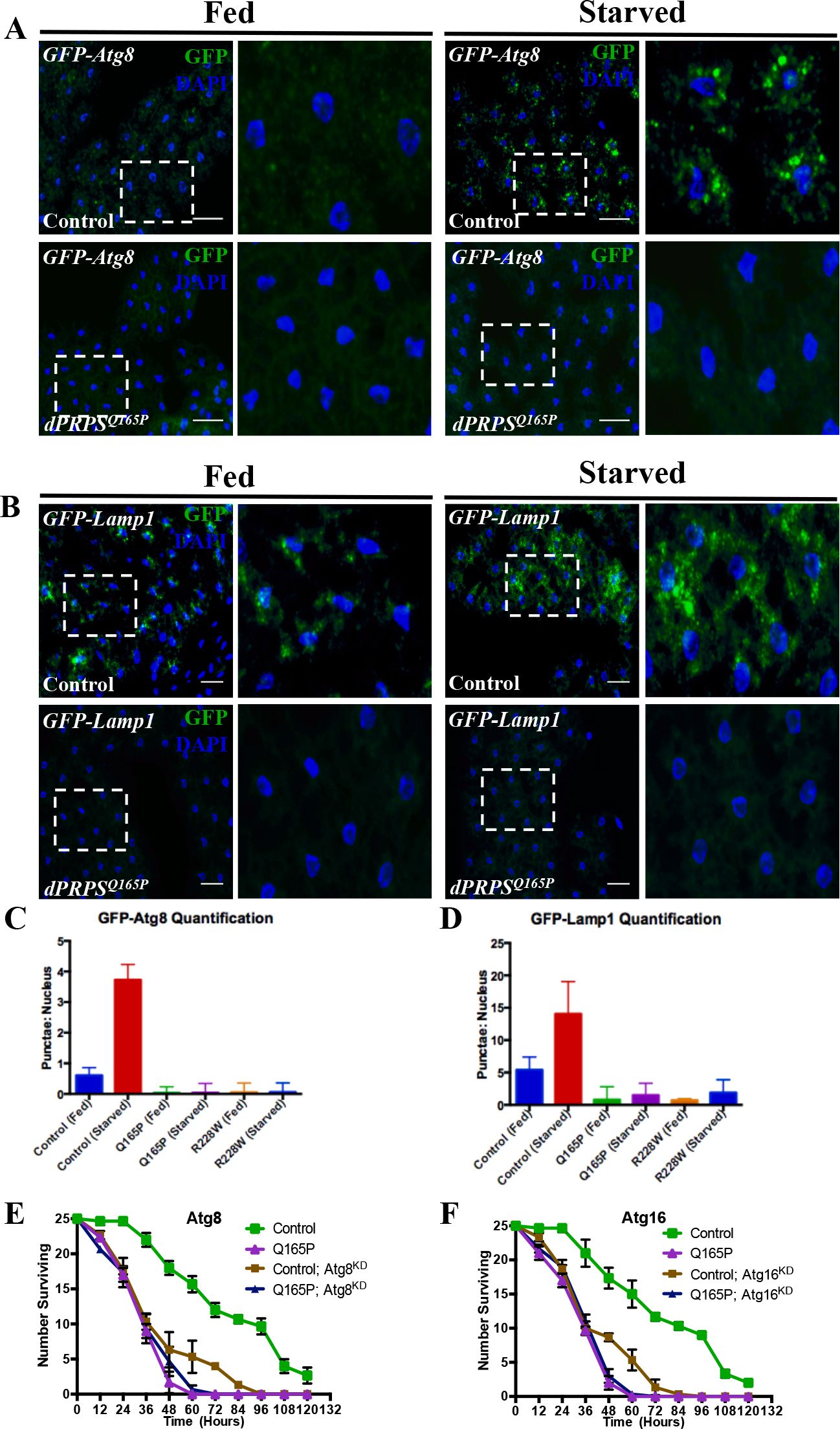
dPRPS is required for starvation-induced autophagy. (A) GFP-tagged Atg8 (GFP-Atg8, green) was expressed via *lsp2-Gal4* to monitor autophagy under fed and starved conditions in adult fat bodies. DAPI is used to visualize nuclei (blue). (B) GFP-tagged Lamp1 (GFP-Lamp1, green), a lysosomal marker, was used as an alternate molecular marker to monitor autophagy. Scale bars: 100 μm. (C and D) Graphs show quantification of GFP-ATG8 and GFP-Lamp1 signals under fed and starved conditions. The number of GFP puncta per nuclei are counted as a marker of autophagy. Error bar indicates standard deviation from triplicated experiments. (E and F) Graphs show survival of wild-type and *dPRPS* mutant flies expressing RNAi against *atg8* (E) and *atg16* (F) via *lsp2-Gal4* under starved conditions. Error bars indicate standard deviation from triplicated sets.

### *dPRPS* mutants have lysosomal dysfunction

The lack of basal Lamp1-GFP signals even under well-fed conditions (Fig. 3B) prompted us to investigate lysosome function in *dPRPS*^*Q165P*^ and *dPRPS*^*R228W*^ flies. To confirm this observation, we directly visualized endogenous Lamp1 proteins using an anti-Lamp1 antibody. Similar to Lamp1-GFP, *dPRPS* mutants have an undetectable level of Lamp1-lysosome puncta under well-fed conditions (Fig. 4A and F4C). Molecular makers for other organelles such as Calnexin, an endoplasmic reticulum protein, and Rab5, an early endosomal marker, showed no visible difference between control and *dPRPS* mutants (Fig. 4A). These results suggest that the lysosome is specifically affected when the function of dPRPS is compromised. We also used LysoTracker, a membrane-permeable dye marking acidic compartments, to visualize the lysosome. Supporting the notion that lysosome function is severely compromised, no LysoTracker signal was observed in the *dPRPS*^*Q165P*^ and *dPRPS*^*R228W*^ fat bodies while readily detected in control (Fig. 4B lower panel and F4D). In contrast, MitoTracker signals were detected in both the control and *dPRPS* mutant fat bodies (Fig. 4B upper panel), indicating that lysosome function is specifically affected by dPRPS hypofunction.

**Figure 4.**
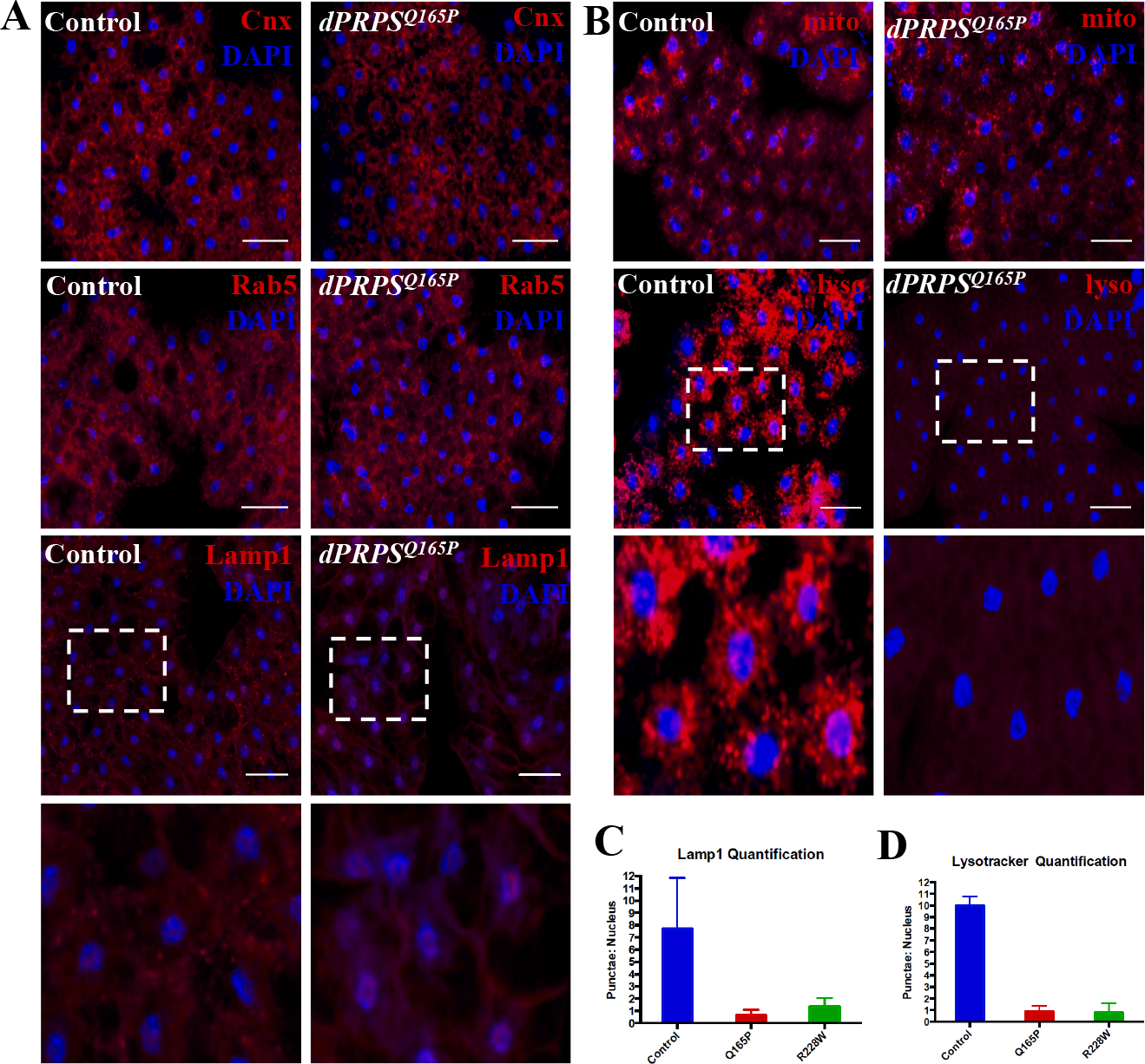
dPRPS mutants have lysosomal dysfunction. (A) Adult fat bodies of control and *dPRPS* mutant flies under fed condition are stained with anti-Calnexin, anti-Rab5 or anti-Lamp1 to visualize the endoplasmic reticulum, early endosome, and lysosome respectively. They are also stained with DAPI to visualize nuclei (blue). For the anti-Lamp1 staining, a magnified view of the indicated area (white box) is presented below. Scale bars: 85 μM. (B) Mitotracker (Mito) and lysotracker (Lyso) are used to monitor the activities of the mitochondria and lysosomes. For the lysotracker staining, a magnified view of the indicated area (white box) is presented below. Scale bars: 70 μM. (C and D) Graphs showing quantification of Lamp1 puncta and lysotracker are also presented.

### *dPRPS* plays a critical function during cellular response to oxidative stress

Autophagy and thus proper lysosome function are needed to protect cells against oxidative stress by removing damaged organelles and molecules (Mizushima 2007). Interestingly, *dPRPS* was identified as one of the transcripts whose expression is induced by oxidative stress (Jin et al. 2015). Indeed, RT-qPCR confirmed that *dPRPS* RNA is upregulated in response to paraquat (PQT) treatment, a Parkinsonian toxin that can induce oxidative stress (Supplemental Fig. S3). We also took advantage of the *dPRPS GFP-trap* allele to monitor dPRPS expression in specific tissues. GFP expression is indeed increased in the ovary of PQT-treated flies supporting the RT-qPCR result (Fig. 5A). However, in the fat body where the basal level of the dPRPS is high, PQT treatment did not have any effect (Fig. 5B). Nevertheless, our results indicate that dPRPS expression is regulated by the cellular level of reactive oxygen species (ROS) and suggests a critical function during oxidative stress. To test if dPRPS is needed for protection against oxidative stress, we examined the survival of *dPRPS*^*Q165P*^ and *dPRPS*^*R228W*^ flies treated with chloroquine, hydrogen peroxide, or PQT under fed conditions. *dPRPS* mutants died faster than control flies when treated with each compound, indicating that they are indeed hypersensitive to oxidative stress (Fig. 5C-5E). We also tested if the hypersensitivity of *dPRPS* mutants to oxidative stress contributes to their susceptibility to starvation. Control, *dPRPS*^*Q165P*^ and *dPRPS*^*R228W*^ flies were treated with NAC under starved condition. While NAC supplementation had little to no effect on control flies, the survival of *dPRPS* mutants were partially rescued (Fig. 5F), suggesting that the oxidative stress indeed contributes to the hypersensitive of *dPRPS*^*Q165P*^ and *dPRPS*^*R228W*^ flies to starvation. Overall, these results suggest that dPRPS play a critical function in cellular response to oxidative stress.

**Figure 5.**
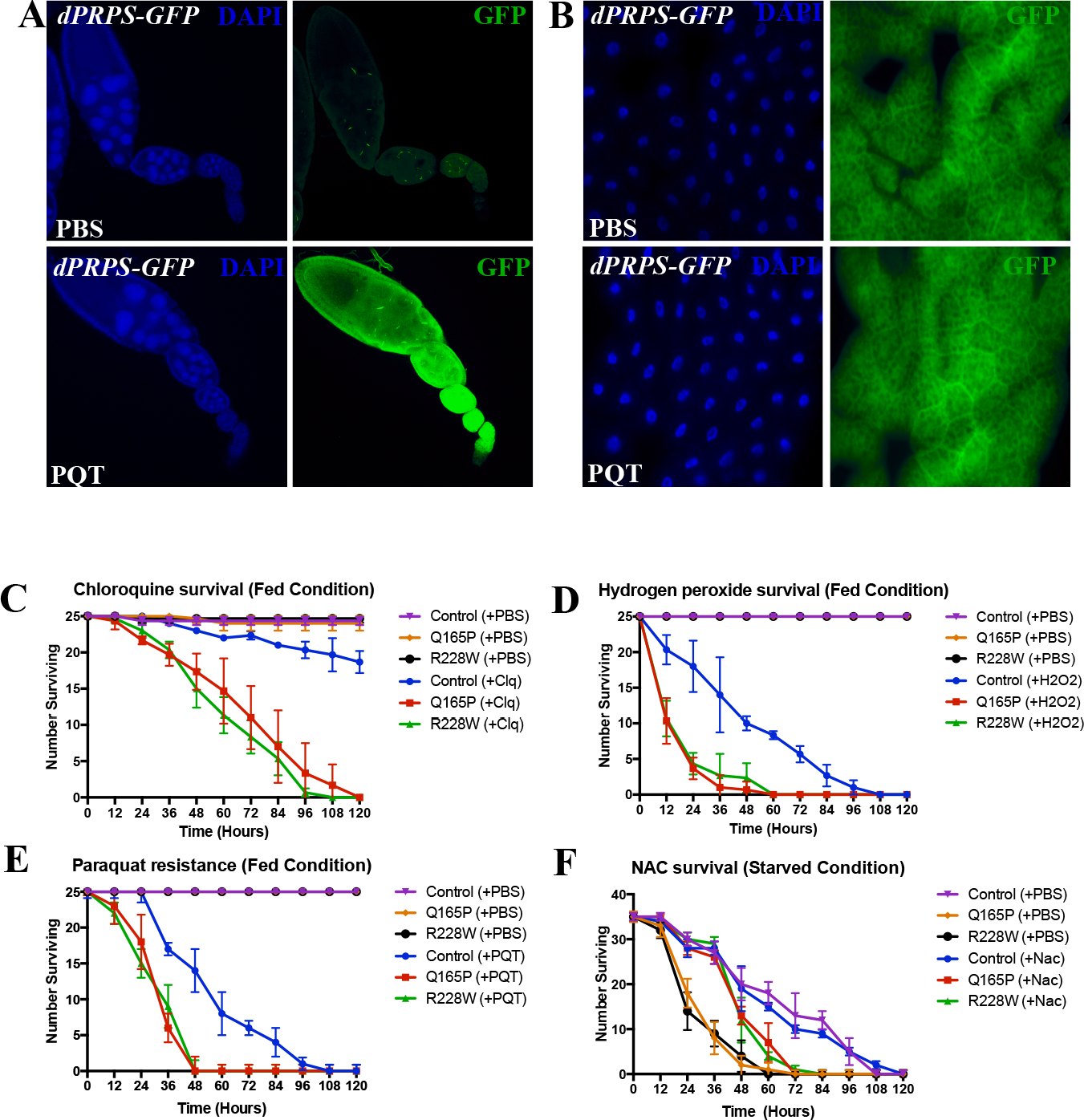
*dPRPS* plays a critical function during cellular response to oxidative stress. (A and B) Ovaries (A) or fat bodies (B) of adult *dPRPS GFP-trap* flies fed with PBS or PQT are immunostained for GFP (green) and DAPI (blue). (C-E) Control and *dPRPS* mutant flies were treated with PBS, chloroquine (C), hydrogen peroxide (D) or PQT (E) under fed conditions. The graph showing their survival is shown. Error bars indicate standard deviation. (F) Under complete starvation, control and *dPRPS* mutant flies are treated with either PBS or 10mM of NAC. The graph showing their survival is presented. Error bars indicate standard deviation.

A spectrum of loss-of-function missense mutations affecting varying levels of PRPS1 activity has been associated with X-linked neurological disorders (de Brouwer et al. 2007). Arts syndrome is the most severe form of *PRPS1*-associated disease, and is characterised by infant death, profound sensorineural hearing loss, intellectual disability, hypotonia and ataxia. Our report here shows that flies carrying the Arts syndrome-derived *PRPS1* mutations, *dPRPS*^*Q165P*^ and *dPRPS*^*R228W*^, have defects in starvation-induced autophagy and profound lysosome dysfunction. Given the high degree of sequence homology between human and fly PRPS (Supplemental Fig. S1) and its essential role in nucleotide metabolism (Hove-Jensen et al. 2017), it is probable that patients with *PRPS1*-associated disorders also have similar defects. However, we have yet to pinpoint the cellular process that is affected by dPRPS dysfunction. One interesting hypothesis is that late endosomal events are affected by dPRPS hypofunction since both lysosome homeostasis and autophagy require factors involved in late endosome trafficking. (Vanlandingham and Ceresa 2009; Jacomin et al. 2016; Nakamura and Yoshimori 2017). For example, the late endocytic machinery is involved in active recycling, degradation and transport of components, such as lysosomal hydrolases and membrane proteins, essential for the maintenance of lysosome function (Huotari and Helenius 2011). Identifying the exactly cellular process deregulated in *dPRPS*^*Q165P*^ and *dPRPS*^*R228W*^ flies will be crucial for understanding PRPS biology and how PRPS dysfunction contribute to neurological disorders.

Because PRPS is a late-limiting enzyme involved in the early step in nucleotide metabolism, it is unclear what are the metabolic changes that lead to autophagy/lysosome defects. PRPP, the product of PRPS (Fig. 1A) is a precursor for purines, pyrimidines, as well as pyridines that are important a wide range of metabolic processes. Extensive metabolic profiling and genetic dissection of the dPRPS-dependent pathways will be necessary to determine the relationship between the nucleotide metabolism and the endolysosomal system. Nevertheless, a surprising finding from our study is that dPRPS expression is tightly linked to the cellular ROS level (Fig 5A and B) and its function is critical for the cellar response to oxidative stress (Fig 5C to 5F). It is well known that autophagy and lysosome are important for protecting cells against oxidative stress (Mizushima 2007) and our study demonstrated that dPRPS is needed for proper autophagy and/or lysosome functions. Notably, in the fat body where the dPRPS expression is naturally high during development, the basal level of autophagy is known to be also high (Zhang and Xi 2015). An interesting speculation is that dPRPS acts as a sensor of ROS and may be one of the important factors that control the level of autophagy for proper cellular response to oxidative stress.

To date, no treatment option is available for patients with PRPS1-associated neurological disorders, such as children with Arts syndrome (de Brouwer et al. 2007; Synofzik et al. 2014; Mittal et al. 2015). Many studies have clearly demonstrated that defects in autophagy are involved in the pathologies of various nervous system disorders, such as Alzheimer’s, Parkinson’s and Huntington’s disease (Bahr and Bendiske 2002; Xilouri and Stefanis 2010; Nikoletopoulou et al. 2015). In addition, there is a growing body of evidence suggesting that lysosome-mediated processes are important for neuronal homeostasis (Nikoletopoulou et al. 2015). Identifying the mechanism by which PRPS regulates autophagy and lysosome homeostasis will most likely uncover a novel treatment option for *PRPS1*-associated diseases.

## Materials & Methods

### Fly stocks

*D. melanogaster* stock cultures were maintained at 25°C. All crosses were performed at 25°C with the exception of the *UAS-RAS*^*V12*^ crosses at 18°C. Following stocks were obtained from Bloomington stock center: *PRPS*^*Mi09551*^ (#53132)*, PRPS*^*Mi09551*^-*GFP-trap* (#59305), *UAS-RAS*^*v12*^ (#4847), *UAS-Lipase* (#67142), *UAS-Atg8* (#58309), *UAS-Atg16* (#58244), *Lsp2-Gal4* (#6357) and the *BSC576* deficiency stock (#26827). The GFP-Atg8 and GFP-Lamp1 are generous gifts from Dr. Steve Jean. (Jean et al. 2015). The *cic*^*Q474X*^ mutant allele used in this project has been previously described in (Krivy et al. 2012). *dPRPS*^*Q165P*^ and *dPRPS*^*R228W*^ mutants were generated through homology-directed protocol as previously described in (Gratz et al. 2015). Positive CRISPR lines were confirmed through T7 Endonuclease I assay following the manufacturer’s instructions (NEB MO302S) and sequenced at the McGill University and Genome Quebec Innovation Centre. Sequence alignment is presented in Supplementary Fig. S1D.

Donor DNA for *dPRPS*^*Q165P*^:

TCCTTTGCATCTCTTTCTACGCTGGCCAATGCACAGAGTCGTGCGCCCATCTCGGCCAAATTGGTGGCCAACATGCTGTCCGTTGCTGGAGCGGATCACATCATCACCATGGATCTGCACGCCTCACCGATTCAGGTAAGTCAGCCCATCAACAACATTTGTATATTTATCTTTGATATTAGAGTGATTTCTTATCGTGC

Donor DNA for *dPRPS*^*R228W*^:

TGCGTAATAGATTAATAATGCTATTAATATTTCACTTTAAGTGTCACCTCAATTGCCGATCGACTGAACGTGGAGTTCGCTCTGATACACAAGGAGTGGAGAAGGCCAACGAAGTGGCCTCTATGGTACTGGTGGGTGATGTCAAGGACAAGATTGCCATTCTGGTCGATGACATGGCCGACACATGCGGCACCATTGTG

Guide RNA (gRNA) as follows:

*dPRPS*^*Q165P*^ forward: GTCGCCAACATGCTGTCCGTTGC
*dPRPS*^*Q165P*^ reverse: AAACGCAACGGACAGCATGTTGGC
*dPRPS*^*R228W*^ forward: GTCGCGCAAGAAGGCCAACGAA
*dPRPS*^*R228W*^ reverse: AAACCTTCGTTGGCCTTCTTGCGC

### Starvation assay

For each genotype, triplicate batches of 25 female flies (<36 hours after eclosion) were transferred to vials of either normal cornmeal medium (fed) or water supply only (1% agarose in H_2_O (starved)). Survival rates were determined by counting the number of dead flies diagnosed by lack of a sit-up response every 12-14 hours.

### NAC treatment

10mg/mL dilutions of PBS or NAC was mixed into cornmeal medium with 25 newly eclosed females (<36 hours) per vial. Every 12-14 hours, dead flies were counted. Triplicate sets were performed.

### Chloroquine, paraquat and hydrogen peroxide treatment

Triplicate batches of 25 2-3 day old females were transferred to cornmeal medium containing 20mM of chloroquine (Sigma), 30mM of paraquat (Sigma) or 1% hydrogen peroxide (Sigma). Every 12-14 hours, dead flies were counted. Triplicate sets were performed.

### RNA extraction and cDNA synthesis

RNA was extracted using the RNeasy Mini Kit (Qiagen) according to manufacture specifications. RNA was collected from whole *yw* flies (n = 5) for the measurement of *dPRPS* transcript levels in adult females fed normal yeast medium supplemented with 1X phosphate buffered saline (PBS), N-acetyl cysteine (NAC), or paraquat (24 hours). 500 ng of RNA was reverse transcribed using the DyNamo cDNA synthesis kit (ThermoScientific).

### Reverse transcriptase quantitative PCR

Reverse transcriptase qPCR (RT-qPCR) experiments were done using DyNAmo Flash SYBR Green qPCR kit (ThermoScientific) according to the manufacture specifications. Threshold cycle (CT) was determined using the Bio-Rad CFX Manager software. *rp49* and *β-tubulin* were both used for normalization. Each primer reaction was performed in triplicates and the three biological replicates were averaged. Primers were designed using Primer3 (*Whitehead Institute for Biomedical Research Primer3 shareware,* Frodo.wi.mit.edu/Primer3). The following primers were used for qPCR reactions:

*β-tubulin* forward: ACATCCCGCCCCGTGGTC
*β-tubulin* reverse: AGAAAGCCTTGCGCCTGAACATAG
*dPRPS Exon4* forward: CTTAGCAAGGGGTGATTTGG
*dPRPS Exon4* reverse: CCTTGATCCACTTGAGTACC
*dPRPS-E1-E3* forward: TTCAGCAACTTGGAGACCTG
*dPRPS-E1-E3* reverse: CCATGGTGATGATGTGATCC

### ATP determination

Five female flies (2-3 days old) were homogenized in 100 *μ*l of 6 M guanidine-HCl in extraction buffer (100 mM Tris and 4 mM EDTA, pH 7.8) to inhibit ATPases. Homogenized samples were frozen in liquid nitrogen, followed by boiling for 5 minutes. Samples were centrifuged and diluted with extraction buffer followed by the addition of luminescent solution (Invitrogen). Luminescence was measured on a luminometer (Turner Biosystems). Relative ATP levels were normalized to the total protein concentration determined by Bradford assay.

### Colorimetric quantification of triglycerides (TAG)

Triglyceride levels were measured using a coupled colorimetric enzymatic triglyceride kit following manufacture instructions (Stanbio). 5 adult flies per genotype were used for each sample measurement in triplicates. Protein levels were measured in conjunction with TAG levels using Bradford assay.

### LipidTox Staining

For whole-mount staining of adult fat bodies (< 36 hours after eclosion), were dissected in 1X PBS and fixed with 4% formaldehyde in PBS for 20 minutes at room temperature. The fixed samples were then washed twice with 0.1% triton in 1X PBS (0.1% PBST) for 10 minutes following incubation in 1X DAPI in 0.1% PBST at room temperature for 15 minutes. After washing with 0.1% PBST, samples were incubated in 1:500 HCS LipidTOX Red Neutral Lipid Stain (ThermoFisher H34476). Samples were visualized within 2 hours by *Leica* SP8 point-scanning confocal system on a *Leica* DMI6000B inverted microscope (provided by the Cell Imaging Analysis Network (CIAN) in the Core Facility for Life Sciences at McGill University).

### LysoTracker and Mitotracker staining

Adult fat bodies were dissected in 1X PBS and stained with either LysoTracker Red (1:1000, ThermoFisher L7528) or Mitotracker (1:300, ThermoFisher M7512) for 5 and 30 minutes respectively. Samples were directly visualized using *Zeiss* Axio Imager fluorescent microscope.

### Immunostaining

The following antibodies were used in this study: anti-Calnexin antibody (1:100, Abcam 75801), anti-Lamp1 (1:100, Abcam 30687), anti-Rab5 (1:100, Abcam 18211), and anti-GFP-FITC (1:200, Abcam). Adult fat bodies were dissected in 1X PBS and fixed with 4% formaldehyde for 20 minutes. Fixed samples were washed twice with 0.3% PBST (0.3% Triton-X-100 in PBS) for 10 minutes. Next, samples were incubated with the indicated antibody. Images were taken using *Zeiss* Axio Imager fluorescent microscope.

### Analysis of lipid droplets and puncta in fat bodies

Fat bodies and lipid droplet size were analysed using *ImageJ* software. Quantification of puncta was performed with *ImageJ* using the particle intensity and 3D objects counter plugin (Bolte and Cordelieres 2006).

---

**Supplementary Figure S1.**
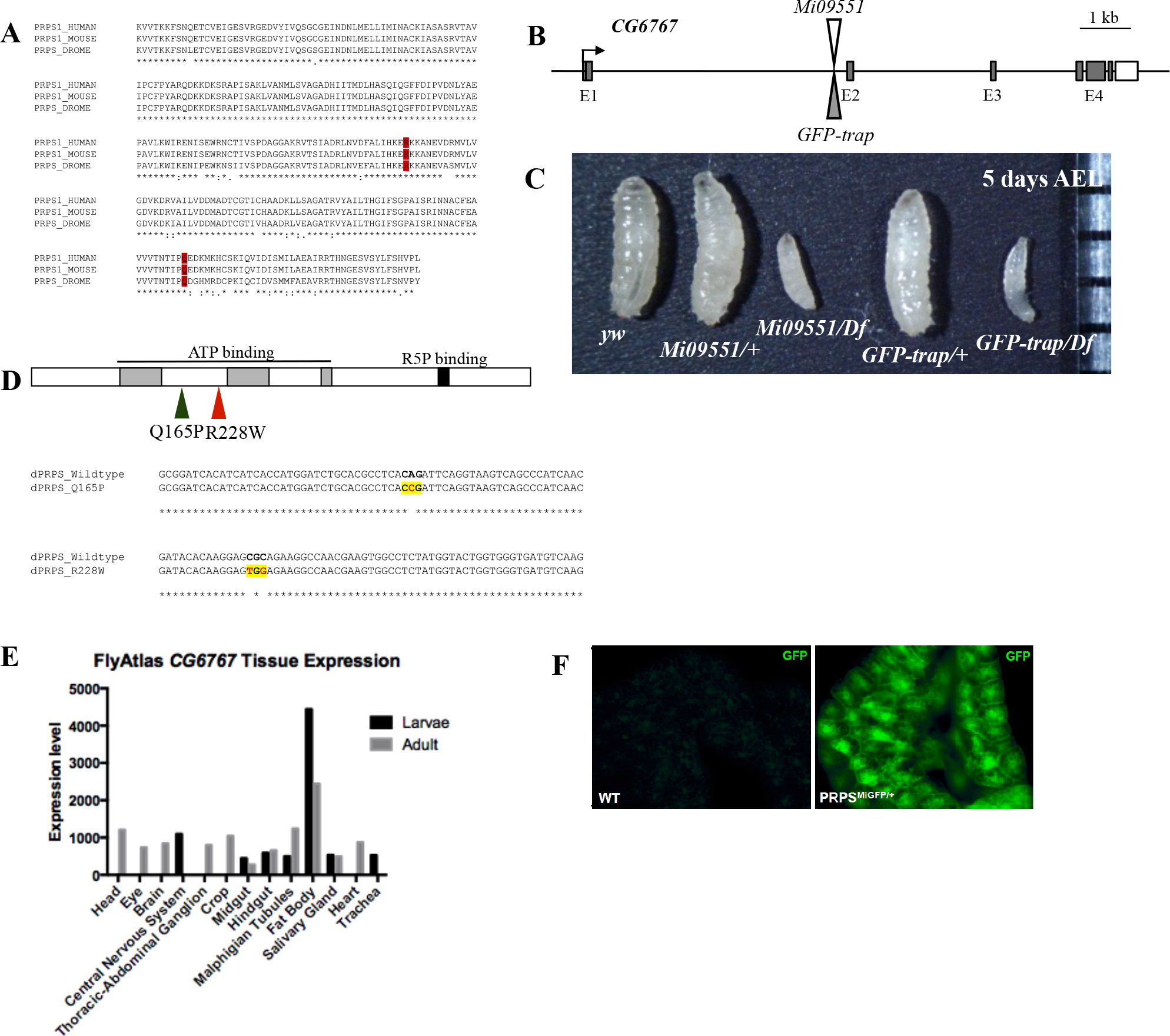
*CG6767* encode the *Drosophila* ortholog of *phosphoribosyl pyrophosphate synthetase* (*prps*) (A) Sequence alignment of human PRPS1, mouse PRPS1, and the only *Drosophila* PRPS ortholog coded by *CG6767* is shown. The *Drosophila* PRPS has 89% protein sequence identity to mammalian PRPS1. Red boxes indicate examples of conserved amino acids that are mutated in human patients with neurological disorders. (B) Schematic of the genomic region of *CG6767* is shown. The two publicly available insertional mutant alleles are marked by arrowheads. Insertions are in the *dPRPS* intron upstream of exon 2. The *Mi09551* allele has a splice acceptor site in the cassette that introduces stop codons downstream of exon 1. The *GFP-trap* allele contains an insertion that inserts the GFP coding sequences in frame at the same location as *Mi09551.* The *GFP-trap* allele produces an internally-tagged GFP-dPRPS fusion protein. (C) The sizes of larvae at five days after egg laying (AEL) are compared. Control (*yw*), *Mi09551* heterozygote (*dPRPS*^*Mi551/+*^), *Mi09551* over a deficiency line (*dPRPS*^*Mi551/Df*^), *GFP-trap* heterozygote (*dPRPS*^*MiGFP/+*^), and *GFP-trap* over a deficiency line (*dPRPS*^*MiGFP/Df*^) are shown. (D) Schematic of patient-derived *dPRPS* mutant alleles, *dPRPS^Q165P^ and dPRPS^R228W^*, are engineered by the CRISPR/Cas9 system. Two conserved amino acids shown in A (red boxes) are edited and the changes in the codon (yellow boxes) are confirmed by sequencing genomic DNA of *dPRPS*^*Q165P*^ and *dPRPS*^*R228W*^ alleles (red letters). (E) Publically available transcript anatomical expression data of *CG6767* from *FlyAtlas Anatomy Microarray* is shown (http://flyatlas.gla.ac.uk/flyatlas/index.htmlmaxdisplayed=30&search=gene&gene=FBgn0036030&radioGene=FBgene). (F) Third instar larval fat bodies of control *yw* (left) and *dPRPS GFP-trap* heterozygous (right) immunostained for GFP are shown.

**Supplementary Figure S2.**
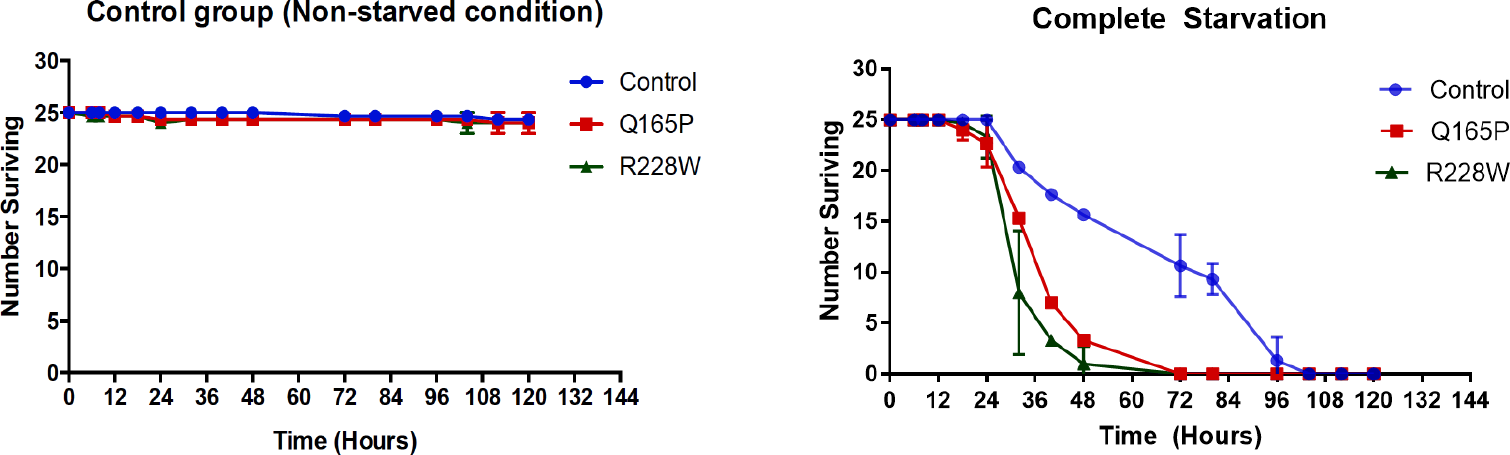
*dPRPS* mutants are sensitive to starvation. Graphs showing survival of control and *dPRPS* mutant adult flies under well-fed (left) and starved (right) are presented.

**Supplementary Figure S3.**
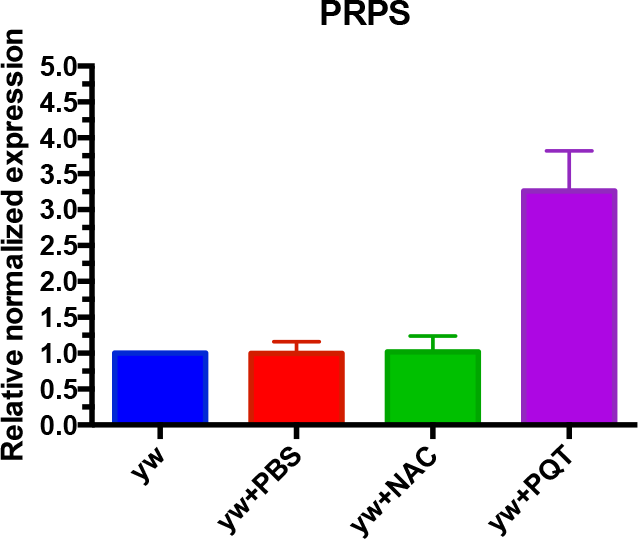
dPRPS expression is induced by oxidative stress. *dPRPS* transcript levels were measured by quantitative RT-PCR. RNA is collected from control flies (*yw*) and flies fed with PBS (PBS), paraquat (PQT) or N-acetyl cysteine (NAC). The graph shows the result from three independent experiments. Error bars indicate standard deviation.

## Acknowledgements

We would like to thank the Bloomington Stock Center for providing fly stocks, as well as CIAN for their assistance in confocal image acquisition. We would also like to thank Dr. Steve Jean for advice and fly reagents. This study was supported by Natural Science and Engineering Research Council of Canada grant 355760-2008 and Canadian Cancer Society Research Institute grant 703339.

## Author contributions

Experiments were designed by NM and performed by KDS and CY. The manuscript was prepared by NM and KDS.

